# Structure and function of the nervous system in the stem of the siphonophore *Nanomia septata*: its role in swimming coordination

**DOI:** 10.1101/2025.11.27.690755

**Authors:** Tigran P. Norekian, Robert W. Meech

## Abstract

The multiple swimming bells, or nectophores, of the colonial hydrozoan *Nanomia septata* are capable of coordinated avoidance swims in both forward and reverse directions. Individual nectophores also contribute to slower forms of swimming during foraging. Communication between a nectophore and the rest of the colony is at cone-shaped structures in the colony’s stem. The stem provides an attachment point for the nectophores and houses the simple nervous system responsible for their coordination. The stem nervous system, revealed by immunocytochemistry, has three main components: two giant axons, a distributed, polygonal nerve network and a set of FMRFamide-immunoreactive nerve tracts. Whereas the nerve network is distributed throughout the stem, the nerve tracts link specific contra-lateral nectophores. Action potentials in the giant axons spread excitation rapidly along the stem, but their connection with individual nectophores is by way of the nerve network. Anatomical evidence is provided for the location of two connecting pathways between the nerve network and the nectophore; one excites an epithelial impulse and leads to reverse swimming; the other provides excitation for forward swimming by feeding into a ganglion-like cluster of nerve cells. Excitation passes to the swimming muscle epithelium by way of a single nerve axon and a nerve ring at the nectophore margin. The work presents physiological evidence for mechanisms, such as facilitation and summation, operating within a multifunctional, bidirectional nerve network, responsible for coordinating epithelial and neural signals in an early-evolved nervous system containing both condensed and distributed units.

**Summary statement:** How a nervous system with two giant axons, a diffuse nerve network and FMRFamide-immunoreactive nerve tracts, coordinates *Nanomia*’s multiple swimming bells to provide the colony with foraging and escape behaviours.

## INTRODUCTION

*Nanomia septata* is a colonial hydrozoan jellyfish with as many as twenty swimming bells, attached in columns on either side of an extended stem (Figure 1A). At rest the whole colony hangs vertically, supported by a gas-filled float (pneumatophore) at its anterior end. The swimming bells, or nectophores, are specialized zooids, which bud off from a growth zone just behind the float (Totton, 1965; Siebert et al., 2015). Together the nectophores are referred to as the nectosome, and below the nectosome is the siphosome with the colony’s other specialized zooids. Individual zooids include hydroid-like gastrozooids, which trap prey using outspread tentacles, bracts, which have a protective function and gonads, all of which are connected to the centrally located stem. The stem houses a simple nervous system, responsible among other things for swimming co-ordination, and the endodermal canal, which distributes the products of digestion.

**Figure 1.**
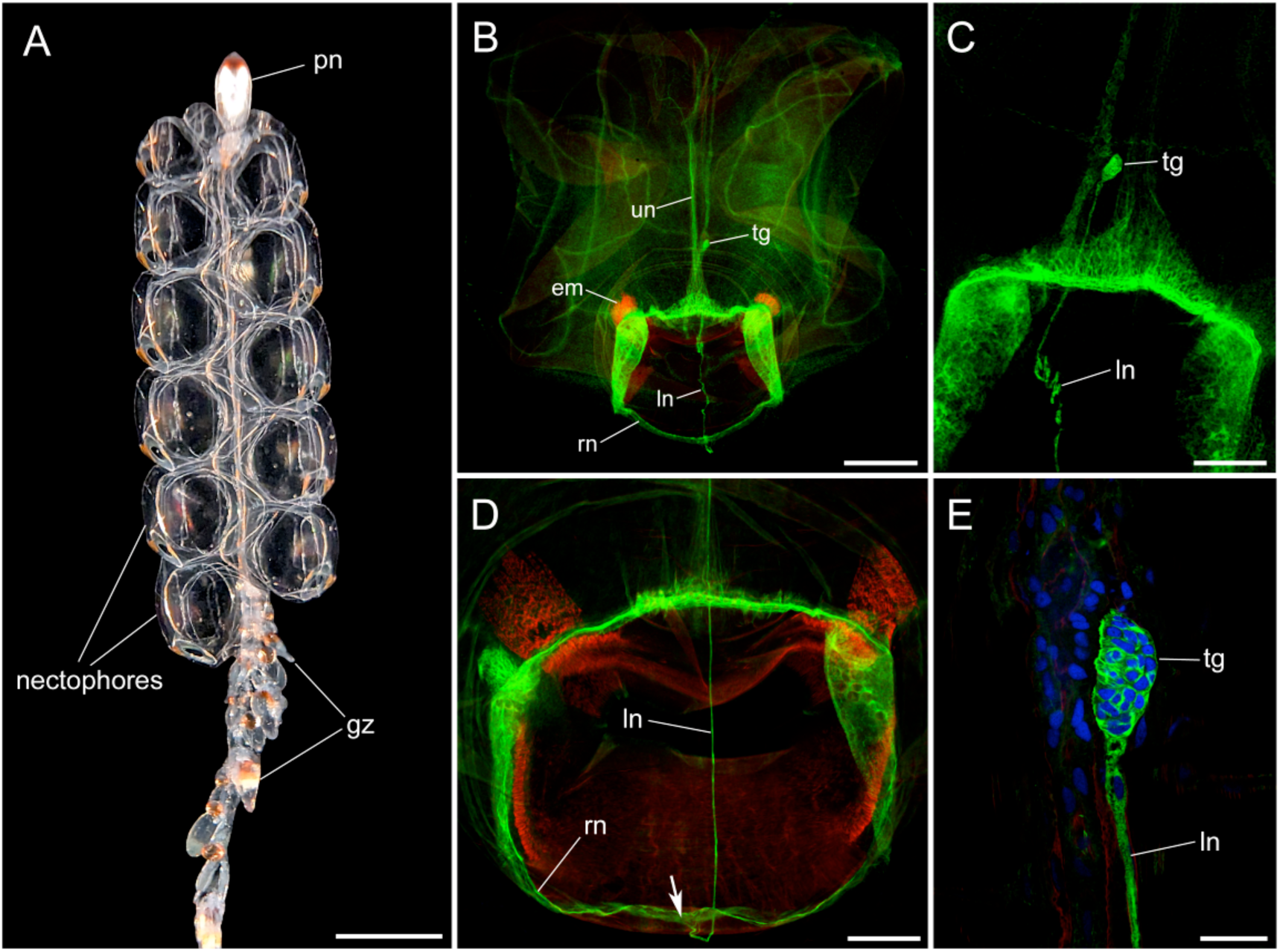
Structure of *Nanomia septata*. (A), colony anterior showing pneumatophore (pn), nectophores and gastrozooids (gz) attached to the central stem, visible through the transparent nectosome. (B), entire isolated nectophore stained with anti-tubulin antibody (green) and phalliodin (red); both the upper nerve (un) and the lower nerve (ln) join the nerve ring (rn); endodermal muscles (em); the terminal ganglion (tg) is located at the back of the nectophore where it normally contacts the stem. (C), the lower nerve (ln) terminates in the terminal ganglion (tg). (D), the point at which the lower nerve (ln) joins the nerve ring (rn) shown by an arrow. (E), terminal ganglion (tg) at high magnification; nuclei (blue) shown by DAPI labeling. Scale bars: A, 5 mm; B, 500 µm; C, D, 200 µm; E, 30 µm;

*Nanomia*’s somewhat ungainly body form is capable of surprisingly varied and agile swimming behaviours. These take the form of asynchronous swims by individual nectophores or blocks of nectophores, and of synchronous swimming by the entire nectosome (Mackie, 1964). *Nanomia* spreads its tentacles or traps food during bouts of asynchronous swimming while coordinated swimming is primarily for avoidance. Coordinated avoidance is of two kinds; a fast reverse swim after a stimulus in the float area and a fast forward swim after a stimulus to the siphosome. The ability to swim abruptly forwards or backwards is powered by contractions in a sheet of epithelial muscle (the myoepithelium) in the bell wall of each nectophore. The direction of travel is set by interactions between the nerves, the muscles and epithelia in the nectophores and in the stem (Mackie, 1964).

A previous publication summarized the neural features of the nectophores (Norekian and Meech, 2020). Now we describe the structure of the nervous system in the stem, its role in swimming coordination and the way individual colonial units show periods of independence and periods of cooperation. We give structural and physiological evidence for the “facilitation barrier” between the stem and the motor neurons in the nectophore and “the blocking or filtering mechanism” regulating the neural traffic between the nectophores and the rest of the colony (Mackie, 1978).

## MATERIALS AND METHODS

### Animals

Adult specimens of *Nanomia septata* (originally *Nanomia bijuga* Delle Chiaje 1844) were collected from surface water at the dock of the Friday Harbor Laboratories, University of Washington, USA, and held in 3 – 4 litre containers at 10°C. Animals survived in good condition for a week or more but wherever possible electrophysiology experiments were carried out on freshly collected specimens. Experiments were carried out in the spring-summer seasons of 2022-2025.

### Immunocytochemistry

In this study we used two markers to identify and study the morphological structure of the nervous system in *Nanomia*. Anti-tubulin immunoreactivity is a well-known and useful tool for identifying many of the neural elements in cnidarian nervous systems. Therefore, as a first marker, rat monoclonal anti-tubulin antibody was used (AbD Serotec, Bio-Rad, Cat# MCA77G, RRID: AB_325003), which recognizes the alpha subunit of α-tubulin, specifically binding tyrosylated α-tubulin (Wehland and Willingham, 1983; Wehland et al., 1983). We have successfully used it to label the neural systems in *Nanomia* nectophores (Norekian and Meech, 2020), the hydrozoan *Aglantha digitale* (Norekian and Moroz, 2020b) and several ctenophore species (Norekian and Moroz, 2016, 2019a, 2019b, 2020a). In addition, FMRFamide (FMRFa)-like antigens have often provided valuable information about the nervous system (Grimmelikhuijzen and Spencer, 1984; Mackie et al., 1985; Satterlie, 2008, 2014; Satterlie and Eichinger, 2014). Anti-FMRFa antibodies are relatively non-specific for FMRFamide and appear to label the entire family of RFamide neuropeptides, distributed in diverse neural systems across the Metazoa (Grimmelikhuijzen & Spencer, 1984; Greenberg et al., 1988). Consequently, we used the anti-FMRFa antibody (rabbit polyclonal; Millipore, Sigma, Cat# AB15348, RRID: AB_805291) not to locate FMRFamide specifically, but as an additional marker for neural elements. Anti-FMRFa antibodies and anti-tubulin antibodies therefore serve as complementary neuronal markers, and we used both to obtain a broad overview of *Nanomia*’s neural morphology.

The muscles in *Nanomia*’s stem contract strongly when placed in fixative, and so we relaxed the tissue by pre-incubation in high Mg^2+^ solution (1 part 0.3M MgCl_2_ and 2 parts of filtered seawater) for 20 minutes prior to fixation. The nectosome was then fixed overnight in 4% paraformaldehyde in 0.1M phosphate-buffered saline (PBS). In this and in all subsequent incubations the temperature was maintained at 4 - 5°C. Following incubation in the high Mg^2+^ solution, the nectosome maintained its natural form during fixation and most of the nectophores remained attached. (Nectophores are capable of autotomy unless treated with care). The preparation was subsequently washed for several hours in PBS.

The fixed animals were dissected according to the task in hand. To investigate the stem; the stem was isolated by removing the nectophores. To investigate the connection between the nectophores and the stem; much of the nectophore bell was removed leaving only the area in direct contact with the stem. This improved the antibody penetration and gave better access to the contact area during the subsequent microscopy.

The dissected tissues were pre-incubated overnight in a blocking solution of 6% goat serum in PBS and then incubated for 48 hours in the primary anti-tubulin antibody diluted with the 6% goat serum solution (final dilution ratio; 1:50). Following a series of PBS washes over a period of 8-12 hours, the tissues were incubated for 24 hours in secondary goat anti-rat IgG antibody: Alexa Fluor 488 conjugated (Molecular Probes, Invitrogen, Cat# A11006, RRID: AB_141373), at a final dilution 1:30. They were then washed for 12 hours in PBS.

In immunocytochemical double-labeling experiments, we used both the anti- α-tubulin antibody and the antibody against FMRFa. After incubating in the blocking solution (6% goat serum in PBS), the tissues were placed for 48 hours in the anti-FMRFa antibody diluted with 6% goat serum (final dilution 1:200). The anti-tubulin antibody was added to the same solution. After several PBS rinses over a period of 8-12 hours, the tissues were placed for 24 hours in secondary goat anti-rabbit IgG antibody, Alexa Fluor 568 (Thermo-Fisher, Cat# A-11011, RRID: AB_143157), at a final dilution of 1:60. For the anti-tubulin antibody we used goat anti-rat IgG antibody (Alexa Fluor 488 conjugated) in the same solution. After 12 hours of washing, the tissue was mounted on glass microscope slides. As a control, we omitted either the anti-FMRFa antibody or the secondary antibody from the protocol; no labeling was detected in either case.

To label the muscle fibers, we used the well-known marker phalloidin (Alexa Fluor 488 phalloidin from Molecular Probes), which binds to F-actin (Wulf et al., 1979). After running the immunocytochemistry protocol, the tissues were incubated in phalloidin solution in PBS for 8 hours at a final dilution 1:80 and then washed in several PBS rinses for 12 hours. The tissues were mounted on glass microscope slides in Vectashield mounting medium. The preparations were viewed and photographed using a Nikon research microscope Eclipse E800 with epi-fluorescence using standard TRITC and FITC filters, and a Nikon C1 Laser Scanning confocal microscope.

The fluorescent dye 4’,6-diamidino-2-phenylindole (DAPI) was used to label cell nuclei; DAPI binds to adenine-thymine (A-T) rich regions of double-stranded DNA and is blue in UV light.

### Electrophysiology

A fire polished glass suction pipette (0.32 mm OD; 0.22 mm ID) was used to record electrical currents from the stem of *Nanomia*. For stem recording the tip of the pipette was introduced between two nectophores. Providing that the tip was advanced slowly, autotomy of the nectophores was unusual. Once at the surface of the stem, gentle suction was applied to the pipette housing and a small area of stem wall drawn onto the pipette tip.

Electrical currents were recorded using a custom-made ‘loose patch’ clamp amplifier (Roberts and Almers, 1992). The recorded current amplitude depended on the area of membrane drawn into the pipette and the proportion of current flowing to ground across the glass/stem interface. The voltage drops between the tip of the pipette and the input terminal of the amplifier depended on the current flowing across the stem surface and the resistance of the suction pipette. A bipolar electrode, placed at different sites on the stem or directly on individual nectophores, was used to deliver electrical stimuli.

### Analysis of externally recorded currents

The currents recorded at the stem surface were of two main types: a triphasic spike-like event and a more slowly developing ramp-like signal. Similar triphasic spikes have been recorded externally from other axons using a range of different techniques. Tasaki (1959) gives an example of the membrane current recorded from a short length of squid axon separated from the rest of the axon by two partitions; essentially the same arrangement as that employed here with the wall of the suction pipette providing the barriers at either side. The peak of the action potential, recorded at the same time with an intracellular micropipette, corresponded to the peak negative membrane current. Tasaki (1959) showed that, providing the axon had uniform properties, the membrane current was proportional to the second derivative of the propagating action potential. Experimental evidence to support this contention was supplied by Katz and Schmitt (1940) and Rushton (1937) provided a theoretical basis. Sub-threshold currents may be expected to spread from the stimulating to the recording site in a manner that depends on the space constant of the conducting tissue involved. In a complex three-dimensional structure like the stem, the spread of sub-threshold currents may be modeled using a delta function (Jack et al., 1975).

Contractions in the nectosome are accompanied by contractions in the longitudinal muscles of the stem ectoderm. These may be either fast, “twitch” contractions, initiated by nerve impulses, or slow postural contractions arising from propagating signals in the stem endoderm. The myoepithelium of the stem ectoderm is excitable but its electrical activity does not propagate independently. Instead, depolarizing currents can spread through “bridges” of continuity across the mesogloea to the endoderm (Mackie, 1978). There they can summate, and when suprathreshold, indirectly excite the endodermal epithelium (Mackie, 1976b), eliciting a slowly (0.4 m/s) conducting impulse in the “S system” (Mackie and Carré, 1983). Muscle spikes recorded externally from *Agalma*, another Physonectid siphonophore, have durations of about 40 ms (Mackie, 1978) corresponding to those of intracellularly recorded fast events (Chain, 1979; PhD thesis). These fast events appear on the rising phase of a slow depolarization lasting 1 – 2 seconds, similar to the SD wave recorded by Mackie (1978). Evidence for propagation within the endodermal epithelium is based on intracellular recordings from the stem of *Forskalia edwardsii* (Mackie, 1978). Endodermal impulses are evident in many of our records, frequently associated with ramp-like currents, but we have not examined their role in nectophore swimming in any detail.

### Kinematics

To aid us trace the origins of the different stem current signals we recorded the swimming responses of the *Nanomia* colony to electrical stimulation. Images were collected at 240 frames/second using an Apple iPhone mounted on a Wild microscope using a Tridaptor mount (MSM, Tianxin Building, 46 Meilin Road, Futian District, Shenzhen, Guangdong, People’s Republic of China). This allowed us to store the locations of the stimulating electrode and the recording pipette. The time of the stimulus was provided by triggering a photodiode (located in the field of view) to flash at the same time. Video frames were at 4.17 ms intervals and occasionally a flash covered two frames; in this case the frame with the brightest flash was taken to be the stimulus time. By matching the video with the electrical signal we could distinguish between true current signals and movement artifacts. We could also match the different components of the behavioural response with the electrical signatures of the different stem components. Videos were converted into single frames using “Frame Grabber” (Appcano LLC). For presentation purposes the contrast in selected frames was increased using Pixelmator (Apple Inc., One Apple Park Way, Cupertino, CA 95014).

### Mg^2+^ anaesthesia

Seawater combined with isotonic MgCl_2_ (61.7 g/l) is commonly used to reduce motor responses to external stimuli in invertebrate animals. In physiological studies on *Nanomia* Mackie (1978) used an isotonic MgCl_2_/seawater solution in a ratio of 1:5, reporting that this concentration of MgCl_2_ did not affect impulse conduction in the stem but greatly reduced the amplitude of muscle twitch contractions. Even a 1:15 MgCl_2_: seawater solution “has a perceptible damping effect on twitch activity” (Mackie 1976b). Previously we have found that 10% isotonic MgCl_2_ in seawater was sufficient to suppress the contractions of nectophore striated muscle (Norekian and Meech, 2020) and we have used 1:10 isotonic MgCl_2_: seawater again here. Initially the animals were highly sensitive to mechanical stimulation, giving coordinated avoidance responses, but after about an hour these coordinated responses became less evident. It is possible that this delayed change may be because of diffusion barriers.

### Terminology used to specify the axes of the *Nanomia* colony

We follow Haddock et al. (2005) in using the term ‘anterior’ to describe the end of the colony with the pneumatophore. For isolated nectophores we use the terms ‘upper’ and ‘lower’ in place of ‘dorsal’ and ‘ventral’; ‘dorsal’ and ‘ventral’ being retained as descriptors for the entire colony.

## RESULTS

### General structure

In each *Nanomia* colony the zooids are grouped into two distinct regions; the nectosome which is at the anterior, just behind the float (pneumatophore), and the siphosome (Figure 1A). The stem, which runs the entire length of the colony, not only serves as a physical anchor for individual zooids but also conducts sensory and neural information. The nectosome has two columns of nectophores with those closest to the pneumatophore being the smallest and most recent. As the nectophores form, they displace the more mature bells further down the nectosome.

The role of the nectosome is to propel the colony through the water by jet propulsion. When a nectophore bell contracts, seawater is forced outwards through the opening at the base (the bell margin). The strength and direction of thrust is determined by the muscular velum, a flap of tissue that fringes the nectophore at its margin (Figure 1D; Norekian and Meech 2020). Tension in the velum is regulated by a sheet of circular striated muscle acting in concert with groups of radially oriented smooth muscles. The bell wall consists of two layers of ectoderm, a layer of endoderm and an elastic mesogloea. The ectodermal layer on the inside surface of the bell, the subumbrella, is a contractile sheet of electrically coupled myoepithelial cells; the ectodermal layer on the outer surface is a simple epithelium consisting of non-muscular, electrically excitable, coupled cells.

Nectophores are not only capable of synchronized contractions, driven by neural impulses in the stem, but also generate swims endogenously. Pacemaker neurons, located within an inner ring of nerves that encircle the bell margin (Figure 1B; Norekian and Meech, 2020), are thought to be the source of this activity, as in other hydrozoans (see Satterlie, 2014). On either side of the nerve ring are two column-shaped nerve plexuses, and entering it at the top and bottom are two nerve tracts. The extensively branching upper nerve appears to have a sensory role (Norekian and Meech, 2020), while the lower nerve (Figure 1B, C) is unbranched until it arrives at, and merges with, the nerve ring (Figure 1D). It consists of a single giant axon (Mackie, 1973) and can be traced to the point of contact between the nectophore and the stem, where it terminates in a ganglion of 40–50 tightly packed neural cells (terminal ganglion; Figure 1C, E; Norekian and Meech, 2020).

### Structure of the neural system in the stem as revealed by tubulin immunoreactivity

The tubulin immunoreactivity (IR) of the stem has two main elements i) a pair of giant axons, ii) a polygonal subepithelial nerve network that connects the giant axons with individual nectophores. The giant axons originate near the base of the pneumatophore (Figure 2A) and run on either side of the stem (Figure 2A-E; 3A, B), so that each has a neighbouring column of nectophores. The axons are 40-50 µm in diameter (Figure 2C; 4B) and in some areas their tubulin-immunoreactive (-ir) processes appear to split into separate fibres (Figure 2C) as if each giant axon is derived from the fusion of many finer nerves. Electron microscopy (Mackie, 1973) shows that the axons have elongated nuclei distributed along their length giving the impression of a syncytial system, “in which fibres can touch without necessarily fusing”.

**Figure 2.**
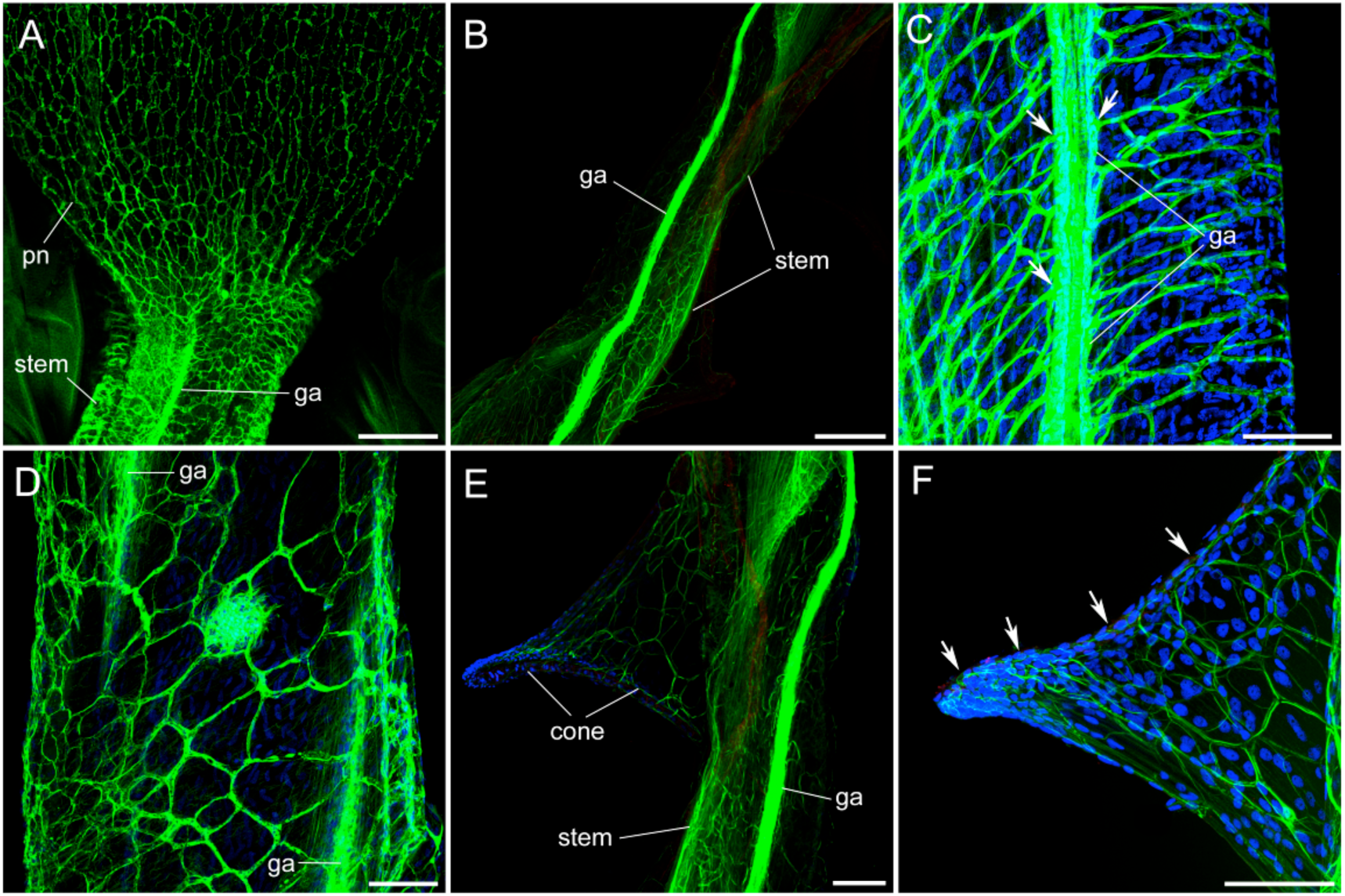
**Stem nervous system labeled with anti-tubulin antibody (green); nuclear DAPI staining (blue)**. (A), stem anterior where it connects to the pneumatophore (pn) showing the origin of the giant axon (ga); the nerve network that covers the outer layer of the pneumatophore merges into the stem. (B), giant axon (ga) running along the entire stem. (C, D), the polygonal nerve network covers the subepithelial layer in the stem and merges with the giant axons (ga) at numerous points (arrows). (E), the cone-shaped structure, which serves as a nectophore docking site, contains the same polygonal nerve network as the stem. (F), higher magnification of the cone and its polygonal nerve network; arrows point to the nectophore attachment surface; cell nuclei (blue). Scale bars: A, B, 200 µm; C–F, 100 µm.

**Figure 3.**
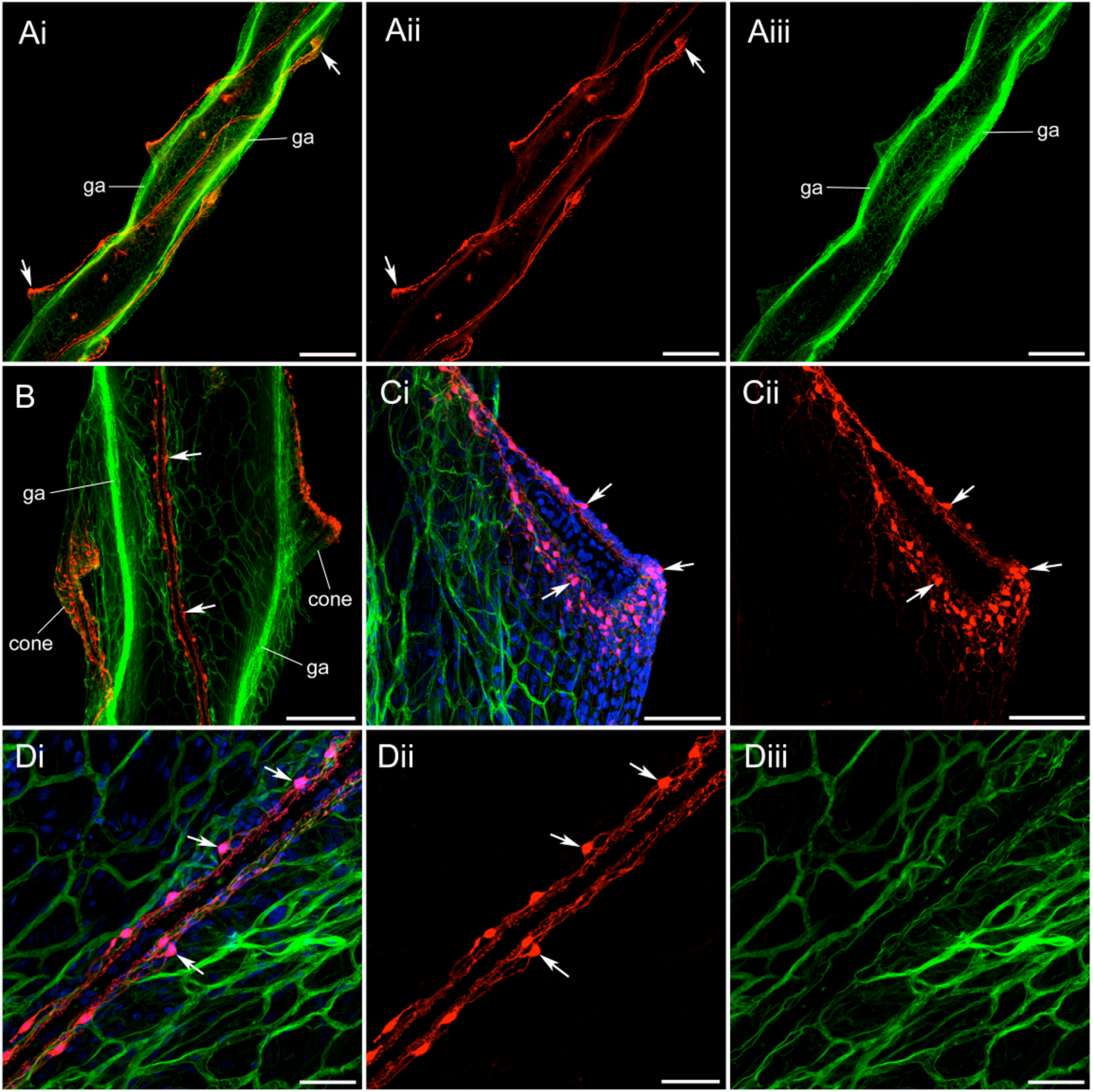
Stem nervous system revealed by anti-tubulin antibody (green); anti-FMRFa antibody (red); nuclear DAPI staining (blue). (Ai), two lateral giant axons (ga) and their associated nerve network (green), together with FMRFa-ir neural tracts connecting contralateral cones. (Aii), red channel of (Ai) showing FMRFa IR only; each neural tract connects contralateral cones (arrows) but not immediate neighbours. (Aiii), green channel of (Ai) showing tubulin IR only. (B), higher power of the stem showing the giant axons (ga), the polygonal nerve network (green), the neural tract (red; arrows), all running along the stem; there is a neural loop at the tip of each cone. (Ci), cone with its polygonal nerve network (green) and FMRFa-ir neural loop (red) at the cone tip. (Cii), red channel of (Ci) showing FMRFa-ir labeling only; arrows show some of the numerous neural cell bodies. (Di), high magnification of the polygonal nerve network (green) and the FMRFa-ir neural tract (red). (Dii), red channel of (Di) showing FMRFa IR only; arrows show some of the immunoreactive cell bodies. (Diii), green channel of (Di) showing that the FMRFa-ir neural tract does not label with tubulin IR. Scale bars: Ai-Aiii, 500 µm; B, 200 µm; Ci, Cii, 100 µm; Di-Diii, 50 µm.

**Figure 4.**
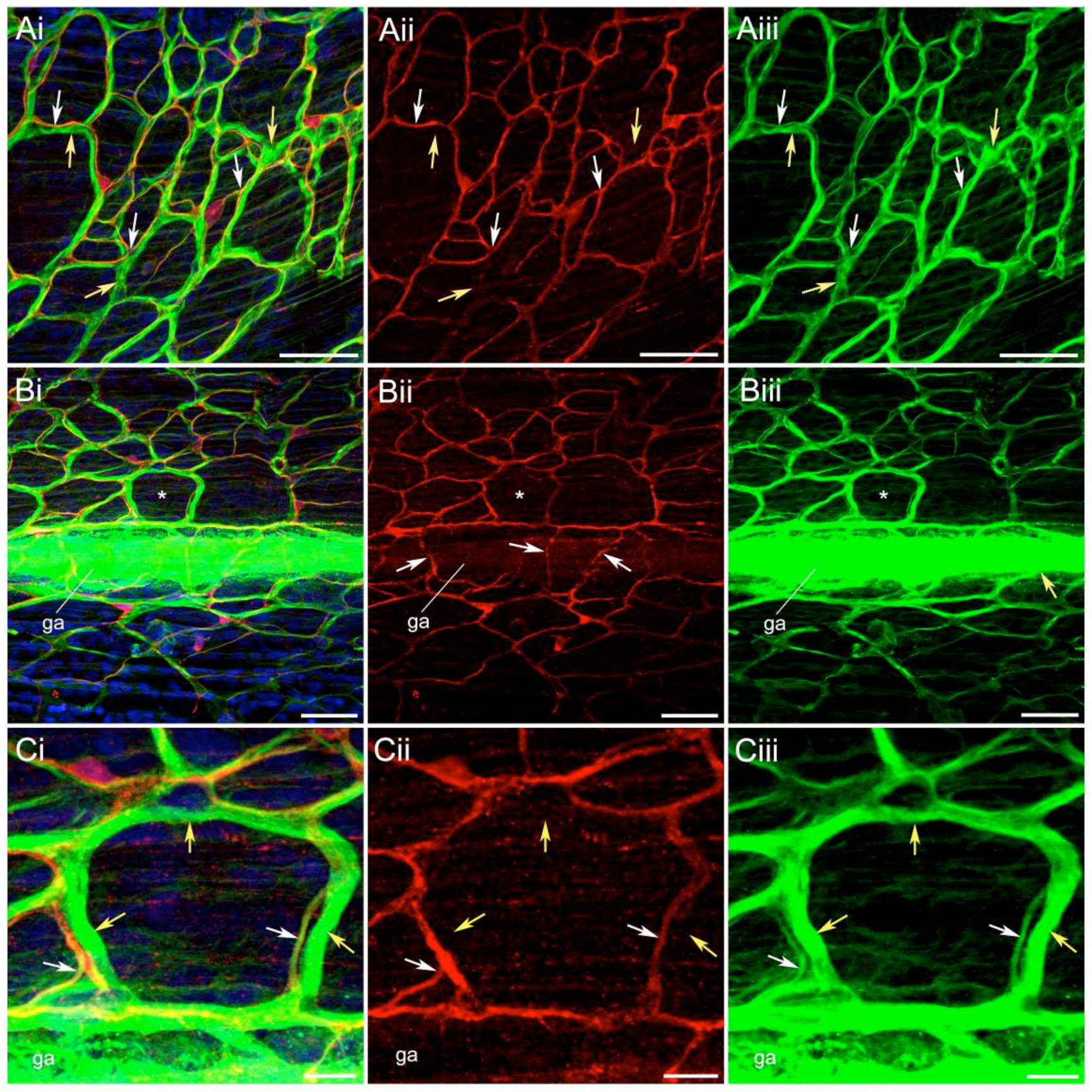
Polygonal nerve network double-labeled with tubulin IR (green) and FMRFa IR (red). (Ai), double labeled image. (Aii), red channel with FMRFa IR only. (Aiii), green channel with tubulin IR only; yellow arrow indicates a thick process with only tubulin IR; white arrows indicate processes with cross-reactivity for both tubulin IR and FMRFa IR. (Bi), double labeled image of the stem with giant axon (ga) and adjacent nerve network; asterisk shows pentagonal shaped neural unit. (Bii), red channel, with FMRFa-ir processes (arrows) that cross the giant axon rather than merging with it. (Biii), green channel showing that the giant axon (ga) is labeled only with tubulin IR; yellow arrow shows tubulin-ir threads merging with the giant axon. (Ci) pentagonal shaped neural unit from Bi; (Cii) red channel with FMRFa IR only; (Ciii) green channel with tubulin IR only. Yellow arrows indicate thick neural threads with tubulin IR only; white arrows indicate processes with both tubulin IR and FMRFa IR. Scale bars: A, B, 50 µm; C, 12 µm.

Merging with the giant axons at multiple places along the stem and located just under the outer epithelium (Figure 2C), is a diffuse, polygonal nerve network, or plexus. Individual units vary in shape from triangular to octagonal and range in size from 10 to 100 µm across (Figure 2D). The network covers the entire stem surface as well as spreading up into the pneumatophore (Figure 2A). It also spreads into the apex of the cone shaped structures shown in Figure 2E, F. These cone-shaped protrusions serve as docking sites for individual nectophores and provide a platform for the neural connection between the central stem and the nectophore.

### Neural elements in the stem revealed by FMRFa immunoreactivity

In the stem of *Nanomia*, FMRFa IR has revealed an additional element of the neural system, not seen with tubulin IR. As Figures 3A, and 6A show, each cone-shaped nectophore attachment point is connected by a FMRFa-ir tract to another cone on the contralateral side - not its immediate contralateral neighbour, but the next one along (Figure 3A; 6A). This double stranded tract has an exceptionally strong signal and does not double-label with tubulin IR (Figure 3B, D). It takes the form of a loop at the top of each cone, on the side contacting the nectophore (Figure 3B, C), and the connecting tract. Both units consist of multiple thin processes and evenly distributed cell bodies (Figure 3B, C, D).

A large part of the polygonal nerve network in the stem, including its thicker threads, has tubulin IR only and shows no FMRFa IR (yellow arrows in Figure 4A). Many thinner processes, however, have both tubulin and FMRFa immunoreactivity (white arrows in Figure 4A). Thus, effectively, there are two nerve networks in the stem. They have matching polygonal structures and are located in the same focal plane (in sections about 30 µm thick). However, while the thick fibers of the tubulin-ir network merge with the giant axons, filaments of the FMRFa-ir network can be seen crossing the giant axons (which have no FMRFa-ir elements) suggesting that they lie slightly below them (Figure 4B). The situation in the cones is similar (Supplemental Figure 1) – part of the polygonal nerve network, including the thicker threads, has tubulin IR only, while part shows double-labeling with FMRFa IR. It is noticeable that the anti-FMRFa antibody stains the neural cell bodies particularly well, unlike the anti-tubulin antibody which labels them rather poorly. Presumably, this reflects the distribution of the antigen molecules concerned.

### The neural connection between the nectophore and the stem

Early developing nectophores, arising at a site near the pneumatophore, are connected to the stem via a thin extended branch (Figure 5A). As new nectophores develop and increase in size, they move along the stem and the connecting branch shrinks. By the time the nectophore is fully mature the connection has been reduced to a cone-shaped structure - the stem cone (Figure 5B, C), described by Mackie (1964) as “a muscular pedicle through which passes an endodermal canal to the subumbrella”.

**Figure 5.**
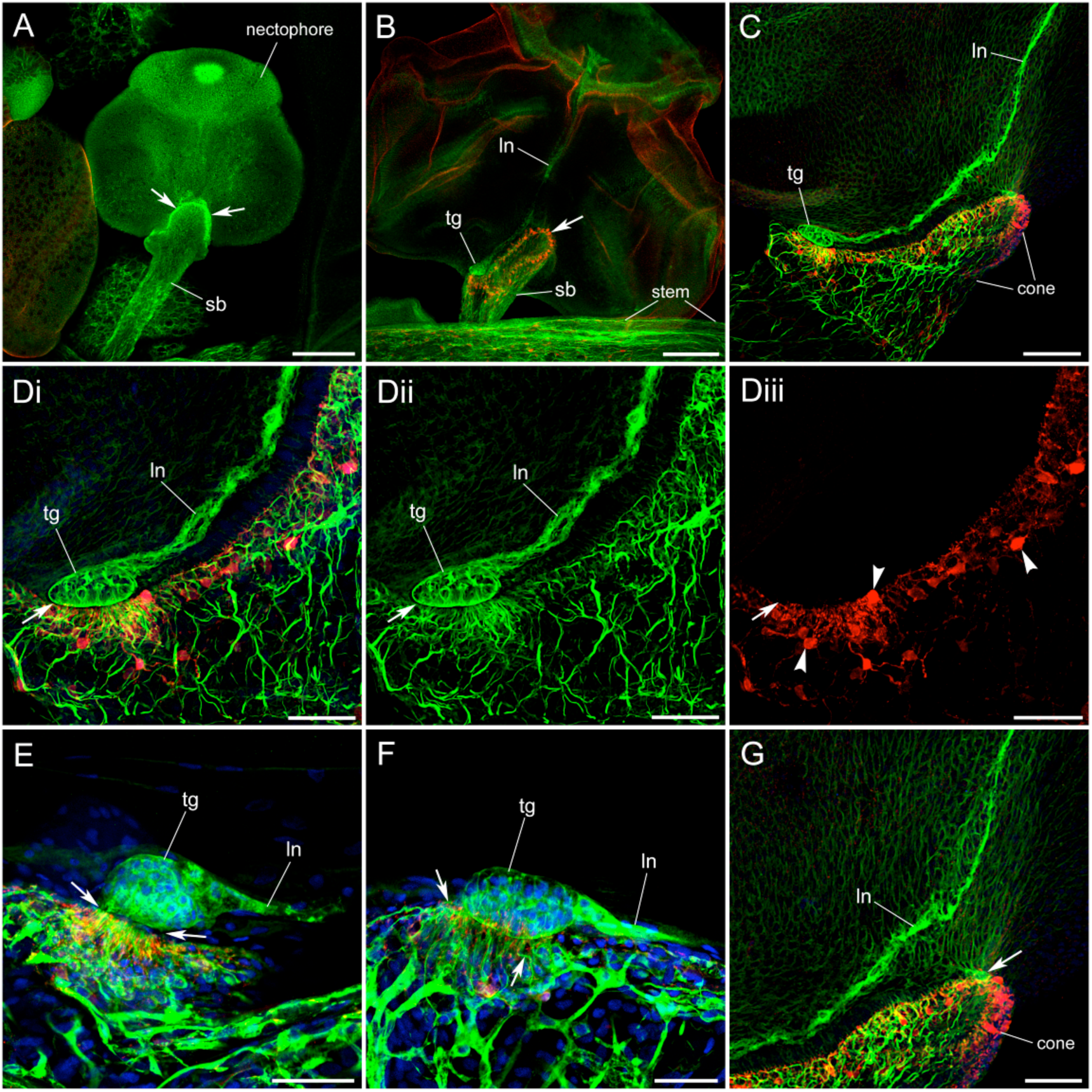
Stem/nectophore connection; anti-tubulin IR (green), anti-FMRFa IR (red), nuclear DAPI staining (blue). (A), early stage nectophore attached to the stem by an extended side-branch (sb). (B), later stage nectophore showing lower nerve (ln) and terminal ganglion (tg) at the point of contact with the shortened stem branch (sb); arrow indicates the FMRFa-ir neural loop (red). (C), mature nectophore connected to the stem via a short cone. (Di), higher magnification of the terminal ganglion (tg) area from (C); the terminal ganglion and cone are separated by a narrow cleft (arrow); many thin nerve fibres, both tubulin-ir and FMRFa-ir, come to the cone surface here. (Dii), green channel of (Di) showing the nerve fibres opposite the nectophore terminal ganglion (tg). (Diii), red channel of (Di) showing FMRFa-ir cell bodies; labeling absent from terminal ganglion or lower nerve. (E), side view of the contact between the terminal ganglion and the cone, showing the thin nerve fibres at the cone surface, opposite the narrow cleft (arrows) that separates the cone from the terminal ganglion; nerve contains both tubulin-ir and FMRFa-ir elements. (F), different view of the terminal ganglion (tg), from the side and slightly above, showing that the polygonal nerve network gives rise to numerous fine processes (arrows) that project toward the terminal ganglion. (G), high magnification of (C) showing the area of contact between the nectophore epithelium and the cone surface (arrow); the outline of the epithelial cells is revealed by background tubulin-ir. D, E and F, different preparations. Scale bars: A, B, 200 µm; C, 100 µm; Di-Diii, G, 50 µm; E, 40 µm; F, 30 µm.

**Figure 6.**
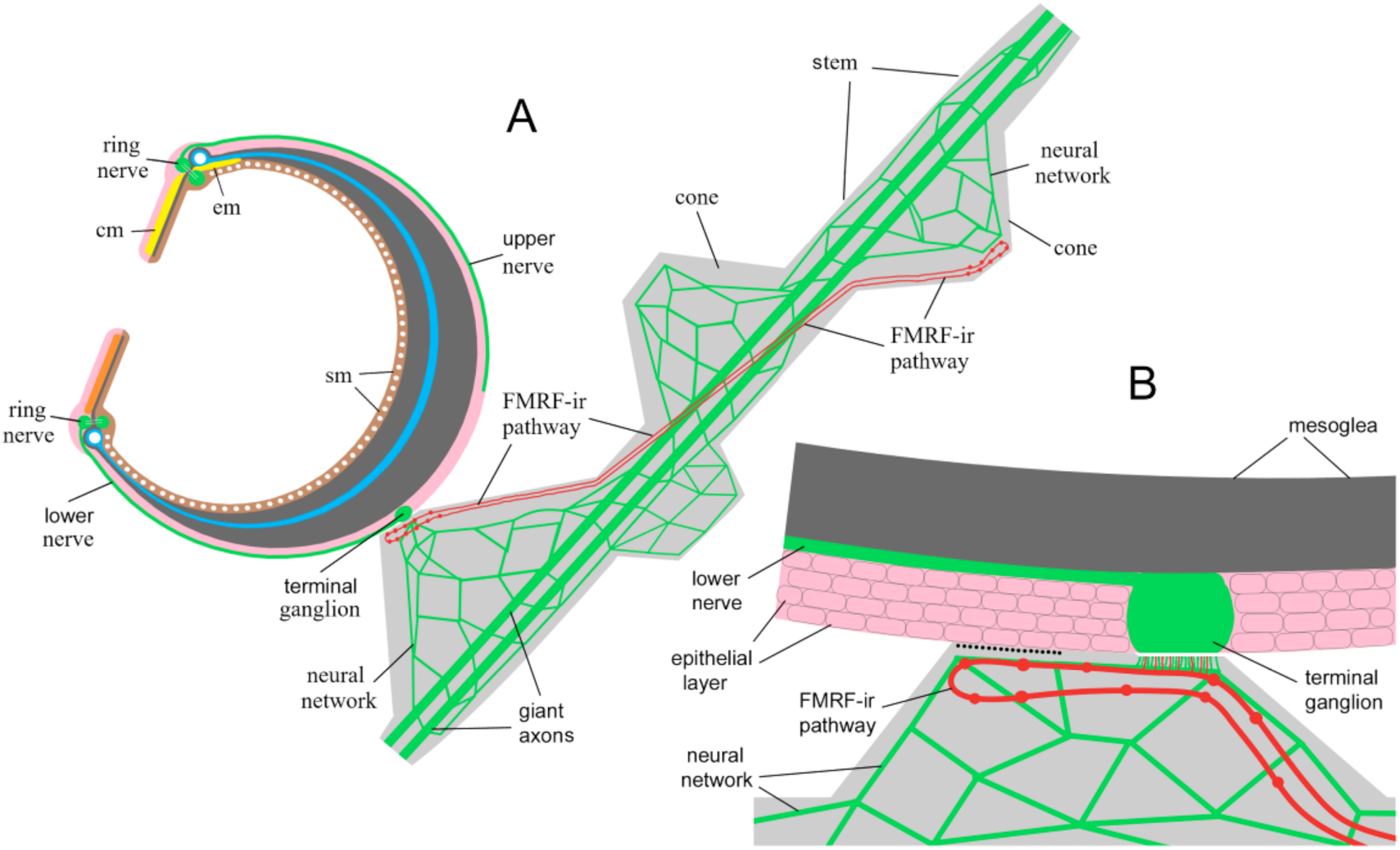
Schematic representation of the neural contact between the nectophore and the stem. (A), the stem neural system includes two giant axons, two tubulin-ir nerve networks and a system of FMRFa-ir, double-threaded, neural tracts between contralateral cones; the stem itself consists of two columns of cone-shaped protrusions that serve as docking stations for the nectophores; the terminal ganglion is located at the attachment point; it is connected to the nerve ring, in the margin of the nectophore, by the lower nerve; also shown are the Claus muscles of the velum (cm) and the muscles of the endoderm (em) that abut onto them (Norekian and Meech, 2020). (B), detailed view of the cone attachment area between the nectophore and the stem; the terminal ganglion of the nectophore is separated from the cone surface by a simple narrow cleft. Outside of the terminal ganglion contact, the epithelial layer of the nectophore is firmly attached to the cone. In this area (black dotted line), electrical junctions may couple the nerve network of the cone with the epithelial conductance pathway in the nectophore.

Mature nectophores communicate with the stem and the rest of the colony via the lower nerve and its terminal ganglion (Figure 5B, C). When nectophores are fixed with the stem in place, the lower nerve is seen to run parallel to the surface of the cone, bypassing almost the entire attachment area before ending in the terminal ganglion (Figure 5D). The ectodermal epithelial layer that covers the nectophore’s outer surface (Figure 5D), is absent at the terminal ganglion itself. Here there is only a narrow cleft between the ganglion and cone surface (Figure 5D, E; 6B). On the stem side of the cleft, many fine nerve fibres approach the cone surface (Figure 5D-F; 6B). These thin fibres arise from thicker threads within the tubulin-ir nerve network and also from the FMRFa-ir double-threaded nerve tract. Outside the terminal ganglion, the epithelial layer of the nectophore is firmly attached to the cone. Here electrical junctions may couple the polygonal nerve network of the cone with the ectodermal epithelial conductance pathway in the nectophore. This epithelial pathway is instrumental in activating Claus’ muscles.

### Physiological function and bell kinematics

#### Control of velum configuration by signals from the stem

An electrical stimulus to the base of the nectosome evokes a synchronized forward swimming response, whereas with anterior stimulation the response has a uniformly backward configuration. Stimuli to the more posterior nectophores sometimes give local responses restricted to the stimulated nectophore or its immediate neighbours. Mid-way along the nectosome, in a “transitional” region, stimulated nectophores may give either forward or reverse synchronized swims (Mackie, 1964).

Figure 7A shows frames from a video of a synchronized swim recorded after stimulating a transitional nectophore. The left-hand frame, collected at the time of the stimulus, shows the numbering system employed for identification. The stimulating electrode touches the dorsal surface of nectophore #4, while the current recording pipette is attached to the stem between nectophores #3 and #4. After 83 ms (centre frame) nectophore #4 had contracted sufficiently for the water leaving the bell to have pushed the velum outwards into an upwardly directed nozzle. The unstimulated nectophores were unaffected at 83 ms, but by 133 ms (right) they were all partly contracted with their vela deflected outwards. The nozzles in nectophores #3 and #4 have been ringed in red and blue and enlargements are shown in the boxes at the far right (cf. Mackie, 1964). Arrowheads show the direction of water flow.

**Figure 7.**
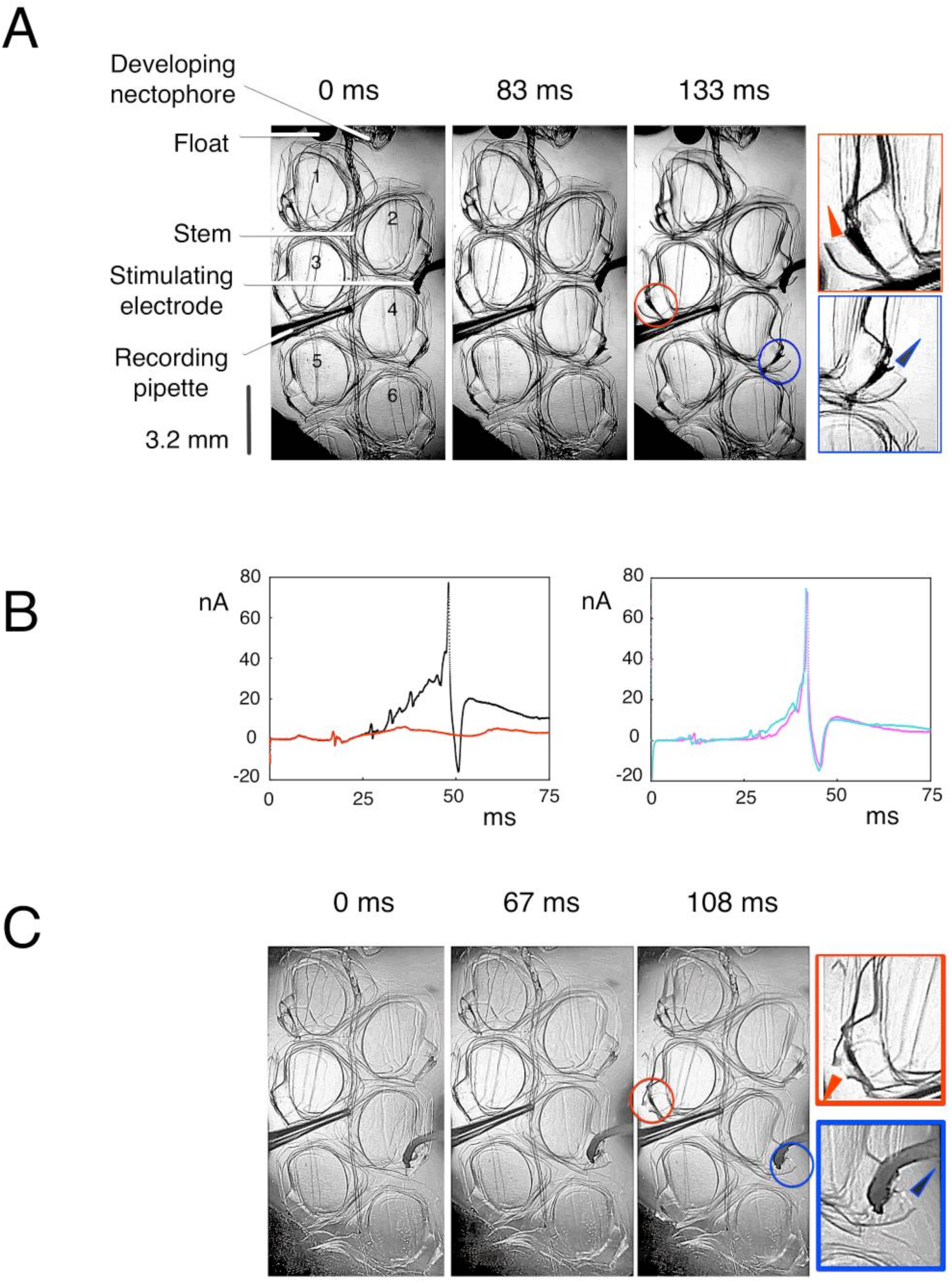
Nectosome kinematics and electrical correlates. (A), nectosome with six nectophores (numbered); stimulating electrode on the upper surface of nectophore #4; recording pipette attached to the stem between nectophores #3 and #4; changes in configuration of the velum at intervals after the stimulus (0, 83 and 133 ms); at far-right frames show the velum at high power. (B), stem current after electrical stimulus to nectophore #4; (left), superimposed current responses to two successive stimuli; after the first stimulus (red trace) the stimulated nectophore contracted alone; after the second stimulus (black trace), all six nectophores contracted as in (A); black trace only visible after it diverges from the red trace; it terminates with an “S” potential; (right), superimposed current responses after stimulating electrode moved to near nerve ring; pink trace corresponds to the swim contraction in (C). (C), stimulated electrode on nectophore #4, near its nerve ring; frames at 0, 67 and 108 ms show the velum configuration at different times; unstimulated nectophores have a forward swimming configuration; high power of velum of nectophores #3 and #4 at far right.

When a velum is viewed head-on the roles of the radial and circular muscle fibres are seen more clearly (Norekian and Meech, 2020). About 38 ms after the stimulus, the circular muscles contract, reducing the diameter of the velar orifice and increasing thrust. Water flow deflects the velum outwards and, in the absence of any radial muscle contraction, it takes the form of a symmetrical truncated cone (Gladfelter, 1972). About 12 ms later, the radially organized Claus fibres contract and the upper velum becomes more rigid. The lower, more flexible part of the velum therefore bulges outwards and deflects the flow of water upwards producing a reverse swim.

The variability of the transitional nectophore’s response to stimulation (Mackie, 1964) lead us to explore the effect of different stimulating positions on the bell surface. Figure 7C shows the result of moving the stimulating electrode to a point on the margin of nectophore #4, near the nerve ring. The velum of the stimulated nectophore exhibited the same upwards configuration, but the vela of the unstimulated nectophores all took up a forward swimming configuration (downwardly directed nozzle). The enlarged side-view of the downwardly directed nozzle of nectophore #3, at the right hand side of Figure 7C, shows that although the lower surface of the velum was markedly contracted, the Claus fibres remained in a relaxed state.

The current flowing in the stem after electrical stimulation of nectophore #4, was recorded using a nearby suction pipette (Figure 7B). The left-hand panel shows the current responses to two successive stimuli. After the first stimulus (red trace) the stimulated nectophore contracted alone, but after the second stimulus (black trace), the six nectophores contracted together. The black trace is only visible after it diverges from the red trace and begins a 30 nA ramp-like climb. The tri-phasic spike at the point of divergence is followed by three further tri-phasic events culminating in a large transient (about 100 nA peak to peak). Following Mackie (1964) we attribute this transient to currents associated with an action potential (“S” potential) in the epithelium lining the endoderm canal (see Methods). The progressive ramp-like current is attributed to the summation of events in the stem muscles (see Mackie, 1964); the muscle epithelium is electrically coupled to the cells of the endodermal canal so that depolarizing events in the muscle summate and depolarize the endoderm to threshold. In the right hand panel of Figure 7B the “S” potential had the same threshold as before, but the ripples and spikes on the rising phase are less distinct. The difference may reflect a small change in the position of the recording pipette.

#### Transfer of excitation from nectophore to stem; the role of the lower nerve and the nerve network

The spread of excitation from the nectophore to the stem was examined by stimulating different sites on the nectophore surface, the current recording site remaining fixed. Approximate stimulating positions are shown in Figure 8A which also shows the path of the lower nerve between the nerve ring and the lateral ganglion (green). Figure 8B shows current records (superimposed) following three successive stimuli with the electrode touching the nectophore’s upper surface (position #6). The early current transients, identified as a ramp (R) and a triphasic spike (S), were recorded before any visible contraction in either the velum or the nectophore (see green bar). A lower amplitude triphasic event riding on a slower deflection was also recorded, as well as a large (20 – 25 nA amplitude) slow wave that peaked at 60 ms. A fast triphasic spike appeared on the falling phase of the slow wave. All three records were associated with a backward swim configuration in the stimulated nectophore.

**Figure 8.**
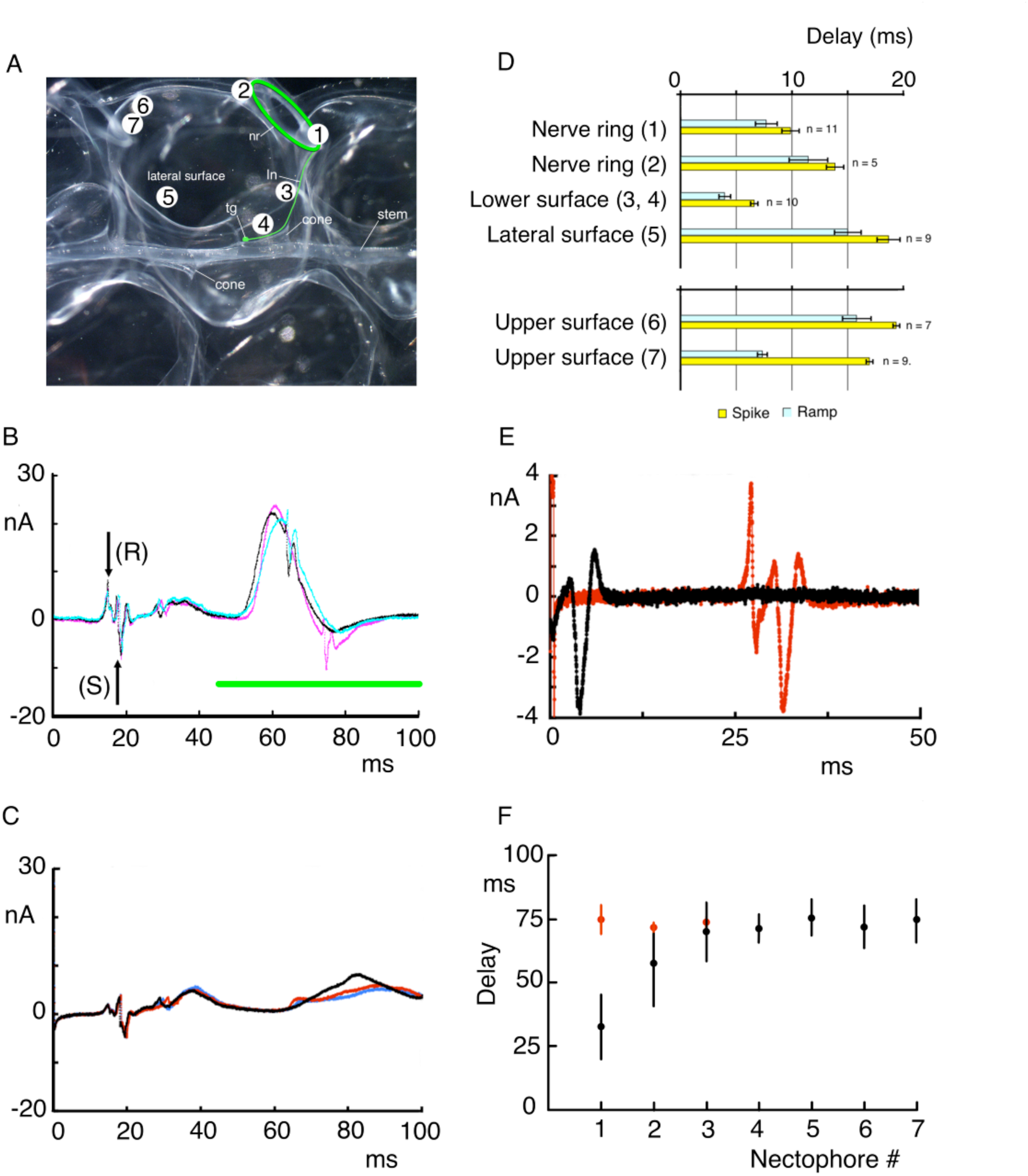
Effect of stimulation site on stem currents. (A), nectophore and stem cone attachment area, showing the pathway (green) of the lower nerve (ln) from nerve ring (rn) to terminal ganglion (tg) and the different stimulation sites (1 – 7). (B), superimposed current records after three successive stimuli to the upper surface (site 6); records have “ramp” (R) and “spike” (S) components; timing of nectophore contractions indicated by green bar; all records associated with a backward swim configuration of the stimulated nectophore. (C), superimposed current records after three successive stimuli to the nectophore lateral surface (site 5); all records associated with a forward swim configuration of the stimulated nectophore. (D), mean delay (+SD) from stimulus to ramp, or to spike, at different locations (1 – 7). (E), superimposed currents after a stem stimulus (black) and a stimulus to a nearby nectophore (red); stem recording site remained unchanged. (F), delay (+/- SD) to just discernible velum contraction response in different nectophores following stimulation of a posterior nectophore (black) or following stimulation to the nearby stem.

When the stimulating position was changed to one nearer the recording electrode on the nectophore’s lateral surface (position #5), the early currents once again consisted of an initial ramp followed by a spike (Figure 8C). The slower ramp with its low amplitude triphasic spike was also present; notably absent however, was the large slow wave. Note that in position #5 all stimuli produced a forward swim configuration in the stimulated nectophore. This confirms preliminary observations by Mackie (1964), which suggested that the upper zones of the nectophore retained the ability to reverse swim (in response to direct stimulation) longer than other regions; forward swimming appearing first in the inferior-lateral areas.

For any given stimulation site, the early currents were highly consistent. Their timing differed, however (Figure 8D). The delay (from stimulus to ramp or from stimulus to spike) was shortest with the stimulating electrode on the lower surface of the nectophore, near the lower nerve and between the terminal ganglion and the nerve ring. The longest delays were when the stimulating electrode was furthest from the lower nerve, on the nectophore’s upper or lateral surfaces.

The delay between the stimulus and the appearance of a current spike in the stem was greatly reduced by stimulating the stem directly. In Figure 8E the simple triphasic current evoked by a stimulus to the stem (black trace) is compared to the combined ramp and spike that followed a stimulus to the nectophore (red trace). The recording site on the stem was the same in each case.

The spread of excitation from a single stimulated nectophore to the rest of the nectosome was assessed from video records of coordinated swims collected at 240 frame/s. Figure 8F shows the delay (+/- SD) between the stimulus and the earliest perceptible velar contraction, in each of the mature nectophores in the nectosome. The black data points show averages (n=7) in which the most posterior nectophore (nectophore #1) was stimulated; in each case the velum took up a forward swim configuration. In the stimulated nectophore, the earliest velar movement depended on the position of the stimulating electrode on the swimming bell (range 16 to 44 ms after the stimulus). The delay from stimulus to the earliest movement in the next-door velum (nectophore #2) was either <42 ms (n=3) or >66 ms (n=4). In nectophores #4 to 7 the mean delay was 73.4 ms (n=27; range 66.6 to 87.5 ms).

Figure 8F also shows the delay to velum contraction when the stem was stimulated directly near nectophore #1; in the case of nectophore #2 the mean delay was 71.9 ms (SD 2.1; n=4). When nectophore #1 was stimulated directly the contraction delay in nectophore #2 (mean, 57.7 ms; SD 16.9) was significantly less than that observed after the stem stimulus (*P*=.036; t stat -2.2) despite the fact that the current spike in the stem was recorded after very little delay (Figure 8E).

#### Stem currents during stimulated and spontaneous swim contractions

Although a suprathreshold electrical stimulus to the stem can elicit a rapidly conducting current spike, the spike is not necessarily accompanied by a coordinated swim movement. Figures 9A, B show currents recorded from the stem, in the nectosome’s transitional zone; the stimulation site was one nectophore diameter (about 3 mm) anterior to the recording site. Following the stimulus artifact (at 0 ms), and a positive-going deflection, there was a triphasic “spike” and a series low amplitude waves (as in Figure 8B, C). The conduction velocity of the spike, estimated from the time to its negative-going peak, was about 330 cm/s. Stimuli that generated a single spike (Figure 9A) failed to elicit any form of nectophore contraction. However, a similar stimulus that generated a double spike (Figure 9B) produced contraction of the nectophore closest to the stimulation site. The second spike was followed by a slow wave with a peak at about 30 ms.

**Figure 9.**
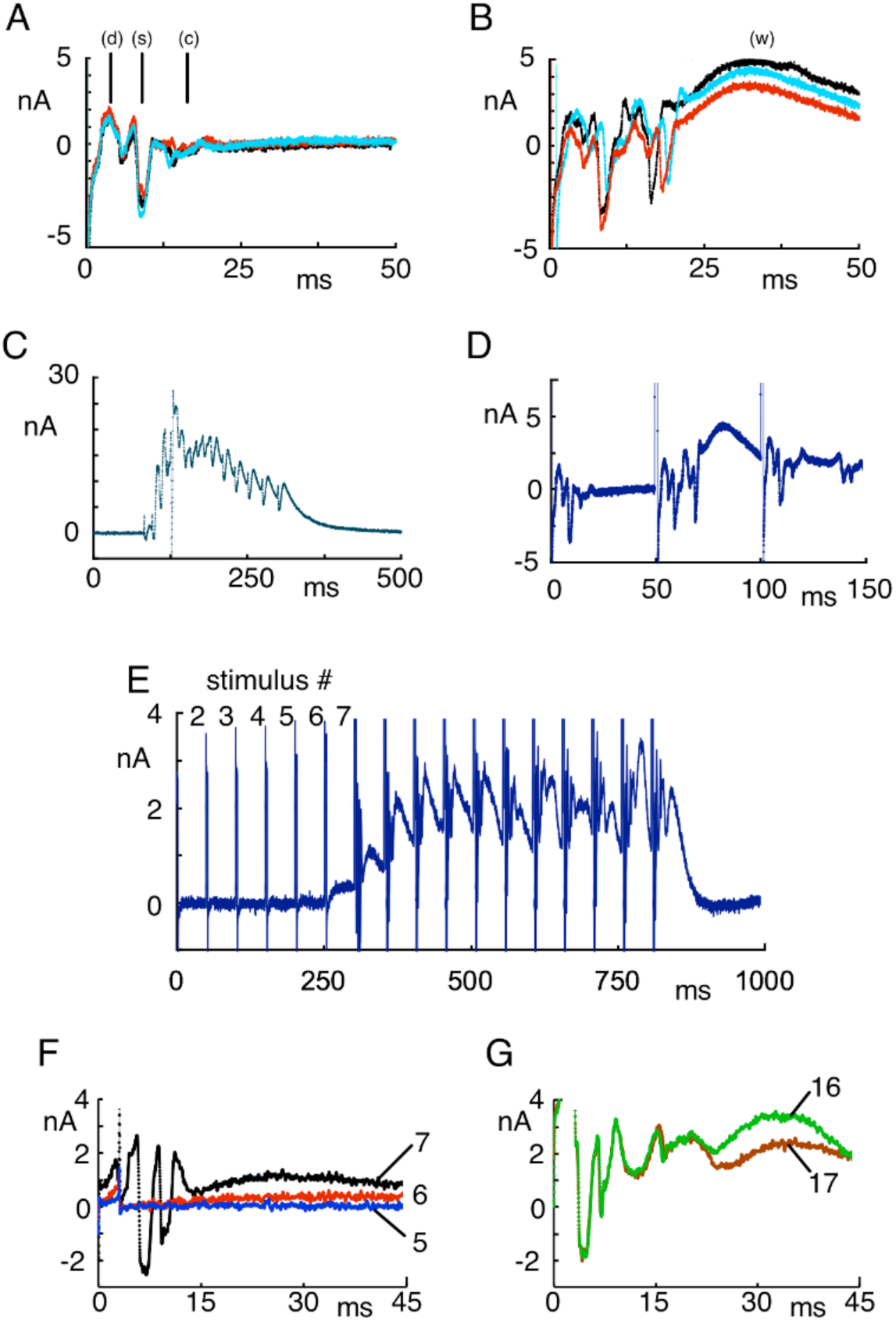
Evoked electrical currents recorded in the stem. (A), stimulus to the stem (artifact at 0 ms) gives a broad deflection (d), a single triphasic “spike” (s), a late complex (c); no swim response; currents from three trials superimposed. (B), as (A) but stimulus followed by two triphasic spikes; there was also a late slow wave (w) and a swim contraction in the nectophore nearest the stimulating electrode. (C), spikes and slow waves during spontaneous swim. (D), repeated stem stimulation, (marked by stimulus artifacts, 50 ms apart); the second stimulus elicited two triphasic spikes and a late slow wave; swim contraction in a nearby nectophore. (E), repeated subthreshold stem stimulation (stimuli 50 ms apart) showing change in excitability. (F), currents associated with stimuli 2 – 4 in (E) were averaged and subtracted from the current responses to stimuli 5, 6 and 7. (G), currents associated with stimuli 2 – 4 were averaged as in (F) and subtracted from the current responses to stimuli 16 and 17.

During spontaneous nectophore contractions, flurries of spikes and slow waves were recorded from the stem (Figure 9C) and the effect could be reproduced by repeated stimuli as shown in Figure 9D. The stimuli were 50 ms apart and the second stimulus elicited a double spike and a contraction in the nearby nectophore. As in Figure 9B the double spike was followed by a slow wave.

#### The role of facilitation in the nectophore stem

Mackie (1964) found that it was possible to evoke asynchronous forward swimming by “prolonged gentle agitation of the siphosome”. This phenomenon is a form of “facilitation” (Satterlie, 2002; Anctil, 2015). Pantin (1933) has suggested that each stimulus leaves behind an after-effect, which could facilitate the transmission of the next stimulus. To examine this possibility we recorded the stem currents during repeated low-level stimulation. In Figure 9E the stimuli were 50 ms apart and there was little visible long-term change in the current baseline until after stimulus #6. For Figure 9F, the currents associated with stimuli #2 to #4 were averaged and subtracted from the current responses to later trials. After stimulus #6 there was a slowly developing positive deflection of the baseline; after stimulus #7 the positive deflection was larger and preceded by a triphasic spike.

## DISCUSSION

Neural, endodermal and muscle systems all contribute to the electrical currents recorded from the stem of *Nanomia* following brief voltage pulses (Mackie 1978). Here we focus on the neural system. Many observations are accounted for by rapidly propagating action potentials in the stem giant axons, but there is also evidence for contributions from a second, separate conducting system. For example, action potentials recorded from the giant axons propagate at velocities of as much as 3 m/s (at 13-16.5 °C; Mackie 1973), whereas Mackie (1964) estimated that during forward swimming, excitation spreads more slowly (0.75-1.50 m/s at 14 °C). Thus, although axon impulses can travel rapidly in either direction, *Nanomia*’s behaviour suggests that there may be two separate pathways. This is supported by the observation that a shock to a bract, gastrozooid, palpon or lower nectophore can evoke the full forward swimming response before the tentacles exhibit any visible shortening, which is the normal accompaniment to giant axon activity.

### Forward swimming

During forward swimming, signals spread from the stem to the nectophore nerve ring by travelling along the lower nerve tract (Mackie, 1964; 1973). Transfer of excitation probably occurs in the terminal ganglion, which is where the lower nerve tract originates (Norekian and Meech 2020; Figures 5D, E, F). Once at the nerve ring, excitation spreads to the muscles of the velum (lower radial and circular) and to the circular swimming muscles of the subumbrella. Contraction of the subumbrella muscles generates thrust while the velar muscles mould the velum into a backward pointing nozzle (Figure 7C; red box).

The giant axons are associated with a network of fine nerves as was first shown by Mackie (1964) using phase contrast and electron microscopy. When a giant axon is penetrated with a dye-filled micropipette the dye spreads into the nerve network (Grimmelikhuijzen et al., 1986) and electrical coupling is demonstrated by showing that action potentials can jump across damaged areas of giant axon (Mackie, 1973). This polygonal neural network covers the entire stem surface and was clearly labeled by tubulin antibody in our experiments. The network elements and the giant axons appeared to merge at sites along the whole length of the stem (Figures 3 & 4). The polygonal network itself appeared to consist of two parts, or two networks with matching morphological structure and location; one, labeled with tubulin IR only, had thick threads, the other with thinner threads was double-labeled with both tubulin and FMRFa antibodies (Figure 4). The existence of two networks corresponds to the phase contrast observation by Mackie (1978) that some of the nerve network appeared to be double-stranded.

Whole-mount studies by Grimmelikhuijzen et al., (1986) using a different RFamide antiserum, also revealed an ectodermal network of multipolar neurons. In some instances, the immunoreactivity was also present in a giant axon and it is possible that the antiserum used, was more sensitive or less specific than the one used here. FMRFa-ir and tubulin-ir networks are also a feature of scyphozoan and cubozoan nervous systems (Satterlie and Eichinger, 2014; Eichinger and Satterlie, 2014).

### Reverse swimming

In the laboratory, reverse swimming occurs most commonly when *Nanomia*’s float contacts the air-water interface. Reverse swims are seen even in nectophores with a damaged lower nerve tract, and they depend on signals propagating within the epithelium of the exumbrellar ectoderm (Mackie, 1964). The stem ectoderm does not propagate impulses and so excitation must spread to the nectophore ectoderm via nerves within the stem (Mackie and Carré, 1983). Presumably this occurs at the point of contact between the nectophore and the stem, i.e. at the cone shaped structure shown in Figure 5C. There is a direct connection from the nerve network in the stem to the nerve cells of the terminal ganglion (Figure 5D, E, F), and a more diffuse association between the nerve network and the nectophore’s ectodermal epithelium. This latter site might provide an electrical pathway between the two and future work might search the area for gap junctions or attempt to pass dye from the nerves to the epithelium at this point.

Once excited, the ectodermal impulse travels to Claus fibres on the left and right of the velum, causing them to contract. The contracted Claus fibres hold the upper velum rigid so that when the subumbrellar myothelium contracts and seawater is forced from the bell, the lower velum bulges outwards forming an upwardly pointing nozzle (Figure 7A). It follows that during a reverse swim the radial muscles in the lower velum remain uncontracted i.e. that signaling in the lower nerve tract/nerve ring must be inhibited in some way. Excitation of the Claus fibres is indirect and involves a neural intermediate located within the nerve ring (Norekian and Meech, 2020). A neural intermediate is also required to link the ectodermal impulse to the swimming muscles of the subumbrella.

### Asynchronous swimmihng and the spread of excitation between nectophores

Although gentle, mechanical stimulation of the siphosome can evoke asynchronous forward swimming (Mackie, 1964), it more generally arises spontaneously, and in the absence of any identifiable external stimulus. Isolated nectophores exhibit long periods of repeated contractions and it is likely that spontaneous swims arise from pacemaker activity in the nerve ring.

A single suprathreshold electrical stimulus to the outer surface of a nectophore in the transitional region of the nectosome, will evoke a local response, but the excitation may also spread to neighbouring nectophores. The direct stimulus excites the ectodermal pathway of the stimulated nectophore and sets up the reverse swim configuration in the velum. However, when the excitation spreads to neighbouring nectophores, they take up a forward swimming configuration. It seems unlikely therefore, that the ectodermal pathway is responsible for the spread of excitation to the stem. More likely is that the pathway runs by way of the lower nerve tract to the terminal ganglion, a suggestion supported by the observation that stimulating the nectophore near the lower nerve tract excites current signals in the stem with the least delay.

The further spread of excitation from the terminal ganglion and along the nectosome during a synchronous forward swim is likely to be by way of the nerve network. We used video recordings (240 frames/s) to monitor this spread and plotted the delay to just visible velum contraction for each of the individual nectophores in the nectosome. In directly stimulated nectophores the delay from the stimulus to the first detectable velum movement was 16 to 44 ms (Figure 8F). After a slightly longer delay (either <42 ms; n=3 or >66 ms, n=4) velum contraction in the nearest neighbour was just visible. In the posterior nectophores the delay was more consistent (mean 73.4 ms; n=27; range 66.6 to 87.5 ms). We conclude that there is a significant delay in communication between the terminal ganglion and the nerve network but that once summation events within the nerve network have exceeded the relatively high threshold of the giant axons, excitation spreads rapidly through the remaining nectosome.

### “The facilitation barrier”

Mackie (1978) could induce swimming by stimulating the stem twice within an interval of 15-20 ms and he suggested that there was “a facilitation barrier” between the stem and the motor neurons in the nectophore. The ventral ganglion, with its large population of neurons, is the possible site of this facilitation.

Single stimuli generate swims providing that they elicit more than one action potential and sequences of repeated action potentials are not unusual. We explored the characteristics of the stem response to repetitive stimulation using different stimulus frequencies. With a 50 ms stimulus interval the axon reached threshold with stimulus # 7 (Figure 9E). However, earlier subthreshold stimuli elicited local responses that may contribute to the axon’s ability to fire repetitive action potentials. After stimulus # 6, there was a slowly rising positive current that reached a broad peak at about 45 ms; after stimulus # 7 the slowly rising current had an increased amplitude and a shorter time to peak.

Repetitive spiking recorded from the stem following stimulation generally had a frequency of about 6 impulses/second (see Mackie 1976b). Similar low frequency impulses in crustacean axons have been modeled using modified Hodgkin/Huxley equations with the addition of a transient potassium current (called I_A_; Connor et al., 1977). To provide a satisfactory model I_A_ must be inactivated by a maintained low-level depolarisation. A current of this kind in *Nanomia* axons would be inactivated by a local response like that in Figure 9F, and this would clear the way for a supra-threshold response. Such channels have been identified in another hydrozoan axon (Meech and Mackie, 1993).

### Role of Summation

Normally, stimulation of an anterior nectophore produces a reverse swim configuration throughout the nectosome. However, we did observe a change in the response following autotomy of one of the nectophores. It appeared that autotomy had not affected the transmission of excitation down the stem but had interupted the signal for reverse swimming. This lead us to suppose that excitation of the exumbrellar ectoderm (necessary for reverse swimming) is a co-operative process with contributions from each nectophore in turn. If the speed of propagation along the stem is high, much of the nerve network would be excited together and be able to exert a cooperative influence. The characteristics of current spread in a multi-dimensional system, such as an epithelium or a syncytium, are described by Jack et al., (1983), who refer to the current “dilution” effect produced by the increased membrane surface in these tissues as compared with a single axon. Any reduction in total current resulting from the loss of a nectophore by autotomy, is likely to cause a rapid spatial decay of current and cause the ectoderm to fail to reach threshold (see Tomita, 1967).

This kind of explanation can account for the finding of Sutherland et al., (2019b), who observed that in freely swimming *Nanomia* the velum can, on occasion, switch from a forward to a reverse swim configuration, mid-way through a swim. We have confirmed this in an electrically stimulated nectophore: stimulation of a transitional nectophore (#1) produced a reverse swim in its next-door neighbour (nectophore #3) but a forward swim in its contralateral neighbour (nectophore #2). However, part way through the swim, nectophore #2 switched from a forward to a reverse configuration. If our supposition is correct, i.e. that cooperative activity in the nerve network is necessary for the ectodermal epithelium to reach its firing threshold, a late contribution from nectophore #3’s network might cause the delayed excitation of the ectoderm system in nectophore #2.

It is possible that during forward swimming, excitation travels along the stem at a speed that fails to generate a cooperative response sufficient to activate the ectodermal system. We have observed, however, that a stimulus to a posterior nectophore which produces forward swims in the rest of the nectosome may be followed some 600 to 1200 ms later by a backward swim. It is as if the first impulse can provide a primer for the second, enabling it to activate the ectodermal system. Supplementary Figure 3 provides a hypothetical basis for the exploration of possible cooperative effects, to be reported in a future communication.

#### FMRFamide-labelled network and neural tract

At present we can say very little about the significance of the FMRFamide-labelled network in the stem of *Nanomia*. In other medusae the FMRFa-ir network includes surface sensory cells (cubomedusae, Eichinger & Satterlie, 2014) and putative sensory cells (scyphozoa, Satterlie and Eichinger, 2014) and in *Nanomia* there is an extensive network of FMRFa-ir cells around the nectophore nerve ring, some of which may be sensory (Supplementary Figure 2; see also Grimmelikhuijzen et al., 1986).

The FMRFa-ir neural tract in the stem connects each nectophore cone with the cone of its contralateral neighbour but one. The very precise nature of this organization, which specifically avoids a direct connection between the cones of nearest neighbours, is highly intriguing. It seems likely that its activity influences the pattern of swimming in some way, but whether it promotes or inhibits excitation remains uncertain. We recorded no instance of an electrical stimulus to one nectophore spreading excitation to distant contralateral neighbours without exciting intervening ones and propose that the FMRFa-ir neural tract prevents an asynchronous swim during feeding becoming a synchronous escape swim. There is a precedent for this; swimming in many hydrozoans is inhibited during feeding and in *Aglantha digitale* the nerve circuits involved are FMRFa-ir labeling (Mackie et al., 2003; Mackie and Meech, 2008; Mackie et al., 2012). Thus the FMRFa-ir neural tract may be part of “the blocking or filtering mechanism” which regulates the traffic between individual zooids and the rest of the colony (Mackie, 1978).

## Acknowledgments

We thank the Director and Staff of the Friday Harbor Labs, University of Wasington, USA for providing excellent facilities, including the Nikon Laser Scanning confocal microscope. RWM thanks Claudia Mills, Eric Edsinger and Bill Gilly for their help and support, thoughtful discussions and for the loan of essential items of equipment. RWM also thanks the University of Bristol for its support. For the purpose of open access, the authors have applied a Creative Commons Attribution (CC BY) licence to any Author Accepted Manuscript version arising from this submission. TPN was supported by the National Science Foundation grants (IOS-2341882; 1557923) and the National Institute of Neurological Disorders and Stroke of the National Institutes of Health (Award Number R01NS114491) to Leonid Moroz.

