## Supplemental Figures 1 to 3 for "Structure and function of the nervous system in the stem of the siphonophore *Nanomia septata*: its role in swimming coordination"

**Norekian & Meech Supplementary Material.**

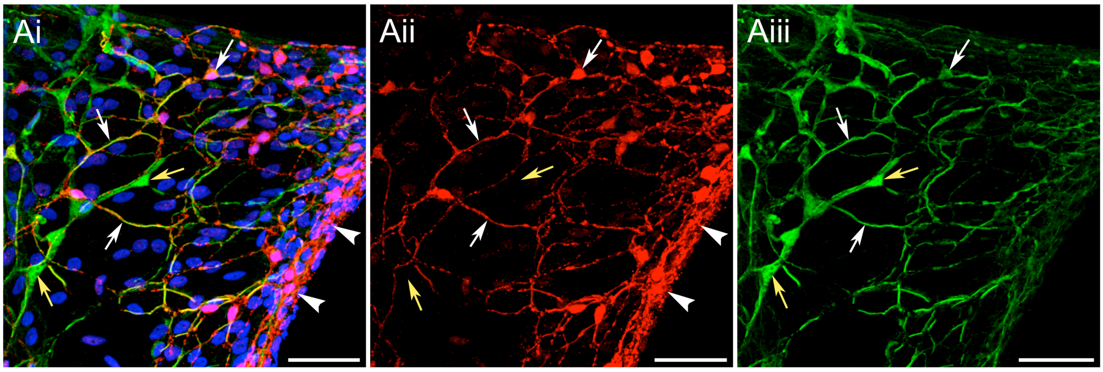


**Supplementary Figure 1: Polygonal nerve network in the cone area.** (Ai), double-labeling for both tubulin IR (green) and FMRFa IR (red) in the cone area; (Aii), red channel with FMRFa IR only. (Aiii), green channel with tubulin IR only; yellow arrow indicates a thick neural processe with only tubulin IR; white arrows indicate processes with cross-reactivity for both tubulin IR and FMRFa IR. Scale bars: 50 µm.

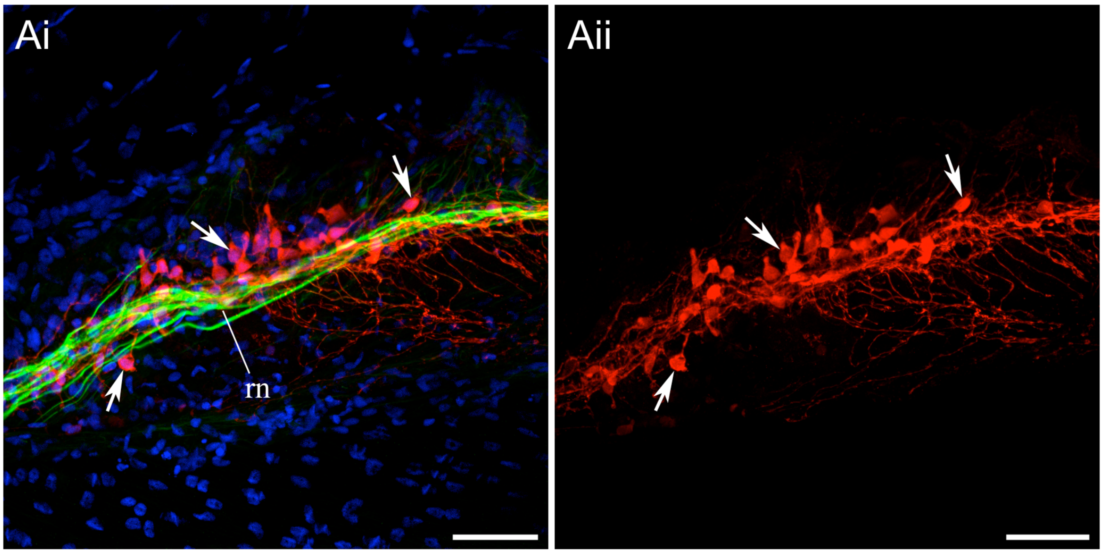


**Supplementary Figure 2: FMRFa-ir neurons in the nectophore nerve ring.** (Ai), nerve ring (rn) near the upper nerve, containing nerves strongly labeled by the anti-tubulin antibody (green) and processes with clearly defined nerve cell bodies (arrows) labeled with anti-FMRF antibody (red). (Aii), red channel showing FMRF-ir processes only. DAPI staining is blue. Scale bars: 50 µm.

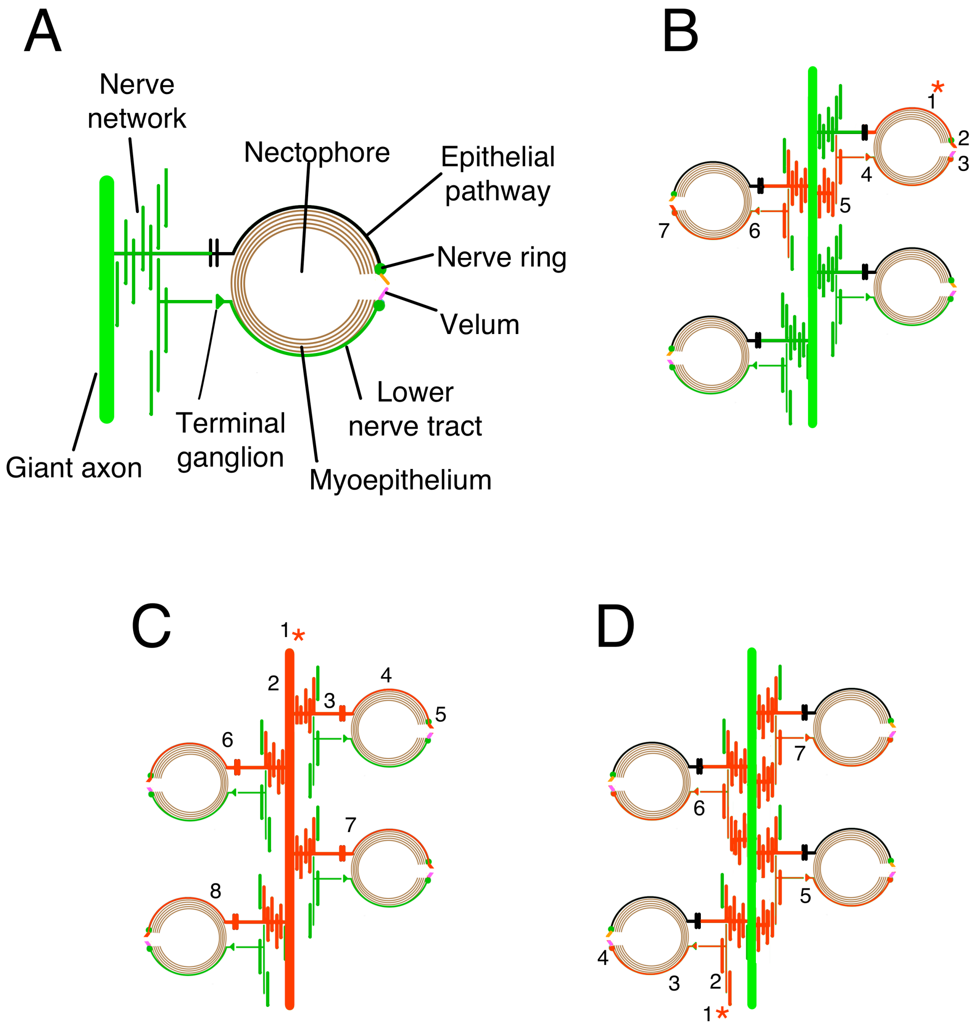


**Supplementary Figure 3: Hypothesis for the coordination and control of *Nanomia* nectophores.** (A) neural and epithelial pathways in schematic form. The pathways include the giant axons in the stem, (for simplicity only one giant axon is shown here) and its associated nerve network. The giant axons are dye and electrically coupled to the polygonal nerve network. The terminal ganglion in the nectophore is taken to be the point of contact between the stem and the nectophore, where excitation spreads from the nerve network in the stem to the lower nerve tract in the nectophore. The nerve network also contacts the ectodermal epithelium of the nectophore near the terminal ganglion and we assume that electrical coupling takes place there. Both the ectodermal epithelium and the lower nerve tract provide an input into the nectophore nerve ring. (There are actually two nerve rings, inner and outer, but for simplicity they have been combined into one). The ectodermal epithelium excites the upper radial muscles, the Claus fibres, of the velum via a neural connection in the nerve ring; the lower nerve tract excites the lower radial muscles of the velum via the nerve ring. There are also excitatory pathways between the nerve ring and the subumbrella myoepithelium of the nectophore. Excited neural units are shown in red throughout.

(B), example of an asynchronous nectophore contraction; here elicited by direct stimulation of the nectophore ectoderm (1*); excitation spreads to the nerve ring (2), and the velum (3), causing contraction of the Claus muscle fibres, but not the lower radial muscle fibres. Excitation from the nerve ring passes to the subumbrella myoepithelium, producing a swim contraction, and also to the lower nerve tract (4). At the terminal ganglion excitation spreads into the stem nerve network (5) and from there it passes to the lower nerve tract of a neighbouring nectophore (6) and the lower radial muscles of its velum (7). Spontaneous asynchronous swims can also arise from pacemaker neurons in the nerve ring and spread to neighbouring nectophores in much the same way.

(C), sequence of events during a reverse swim; excitation of the giant axons, (1*) at the anterior of the animal spreads to the nerve network (2); an impulse in the ectodermal epithelium of the nectophore (4) is then elicited by the spread of current from the nerve network across the electrical synapse (3) at the point of contact between the stem and the nectophore; the epithelial impulse excites the Claus fibres, and the swim muscle of the subumbrella, via neural connections in the nerve ring. Nectophores #6, #7 and #8 are excited to contract in quick succession as the action potentials traverse the nectosome at a velocity of up to 3m/s. [Inhibition may prevent the Claus fibres and the lower radial fibres in the velum being activated at the same time.]

(D), sequence of events during a forward swim; excitation occurs at the rear of the animal – in the siphosome (1*) – and activates the nerve network there (2); excitation spreads to the nectophore via the terminal ganglion and the lower nerve tract (3); at the nerve ring (4), excitation spreads to the lower radial muscles of the velum and the swim muscles of the subumbrella; excitation continues towards the anterior of the animal, propagating in the nerve network and causing the remaining nectophores of the siphosome to contract. The rate of conduction may be slower than for propagation in the giant axons.
